# kTWAS: Integrating kernel-machine with transcriptome-wide association studies improves statistical power and reveals novel genes

**DOI:** 10.1101/2020.06.29.177121

**Authors:** Chen Cao, Devin Kwok, Shannon Edie, Qing Li, Bowei Ding, Pathum Kossinna, Simone Campbell, Jingjing Wu, Matthew Greenberg, Quan Long

## Abstract

The power of genotype-phenotype association mapping studies increases greatly when contributions from multiple variants in a focal region are meaningfully aggregated. Currently, there are two popular categories of variant aggregation methods. Transcriptome-wide association studies (TWAS) represent a category of emerging methods that select variants based on their effect on gene expressions, providing pretrained linear combinations of variants for downstream association mapping. In contrast, kernel methods such as SKAT model genotypic and phenotypic variance using various kernel functions that capture genetic similarity between subjects, allowing non-linear effects to be included. From the perspective of machine learning, these two methods cover two complementary aspects of feature engineering: feature selection/pruning, and feature modeling. Thus far, no thorough comparison has been made between these categories, and no methods exist which incorporate the advantages of TWAS and kernel-based methods. In this work we developed a novel method called kTWAS that applies TWAS-like feature selection to a SKAT-like kernel association test, combining the strengths of both approaches. Through extensive simulations, we demonstrate that kTWAS has higher power than TWAS and multiple SKAT-based protocols, and we identify novel disease-associated genes in WTCCC genotyping array data and MSSNG (Autism) sequence data. The source code for kTWAS and our simulations are available in our GitHub repository (https://github.com/theLongLab/kTWAS).

## Introduction

Transcriptome-wide association studies (TWAS) have emerged as an important technique for associating genetic variants and phenotypic changes[1–5]. Pioneered by Gamazon *et al.*[6], TWAS is typically conducted in two steps: First, a model is trained to predict gene expression from genotypes, using a reference dataset which contains paired expression and genotype data. Techniques including ElasticNet[6], Bayesian sparse linear mixed models (BSLMM)[7–9], deep auto-encoder models[10] and deep learning regression models [11] are used to fit this genotype-expression model. The pretrained genotype-expression model is then used to predict expression activity from the main dataset for genotype-phenotype association mapping (referred to as GWAS dataset hereafter), which contains genotype and phenotype information (but not expression data) for each case or control in the GWAS cohort. As first demonstrated by Gusev *et al.*[8], meta-analysis methods for conducting TWAS using summary statistics from the GWAS dataset have also been developed[12]. The key insight of TWAS is that transcriptomic data can be used to select for genetic variants critical to gene expression (i.e., eQTLs), which improve the quality of downstream GWAS. By modelling the association between linear combinations of variants and expression, TWAS effectively aggregates many genetic variants into a small number of meaningful linear combinations. Remarkably, this approach remains effective even when the predictive power of the genotype-expression model is low. As a result, despite having average *R^2^* values around 1%, the use of genotype-expression models in TWAS has led to significant successes in real data analyses[3, 4, 13–17]. Indeed, as demonstrated in our simulations[18], predicted expressions generated by a genotype-phenotype model can perform better than actual expression data when applying TWAS analysis. This may be because predicted gene expressions capture the genetic component of expressions more precisely than the actual expressions, which include multiple components such as experimental artifacts and environmental factors.

The popularity of TWAS has overshadowed another well-established branch of kernel machine-based models of genetic association, such as the sequence kernel association test, or SKAT[19, 20]. Evidently, with the emergence of TWAS (**Table 1** Column 3,4), the citation of the SKAT paper for common variants analysis is decreasing (**Table 1** Column 2), although the community still cites the SKAT paper for rare variants analysis (**Table 1** Column 1). The key insight of kernel methods is that the similarity of a genetic region between different subjects (as captured by a kernel) can be used to associate genotypic and phenotypic variance in that region, without knowing which specific genetic variants are causal in the focal region. As a result, kernel-based methods can model the aggregated effects of multiple genetic variants and capture genetic interactions within a local region, while being robust to noise. At first glance, TWAS and kernel-based models appear quite different, as TWAS utilizes expressions, whereas kernel methods only use genetic data. Intuitively, TWAS may appear to be more powerful as it integrates more information in the form of expression data. However, in our opinion, TWAS and kernel methods are quite comparable because they are both variant-set analyses which test an aggregated set of genetic variants for associations with a phenotype. Essentially, TWAS selects and weights genetic variants for aggregation using a linear model, whereas kernel methods structure genetic variants into various kernel machines. From the perspective of machine learning, these two methods cover two complementary aspects of feature engineering: feature selection/pruning, and feature modeling. Kernel methods organize genetic variants in a flexible way to cope with unknown genetic architecture, but do not provide a quantitative way to pre-select meaningful variants; while TWAS models select meaningful variants via gene expressions but leave the modeling of them to a simple linear form.

**Table 1.** Number of citations for SKAT and TWAS papers over the last ten years (Google Scholar).

| Year | SKAT(rare)[19] | SKAT(common)[20] | PrediXcan[6] | TWAS(meta)[8] |
| --- | --- | --- | --- | --- |
| 2010 | 1 | **4** | 0 | 0 |
| 2011 | 14 | **32** | 0 | 0 |
| 2012 | 70 | **55** | 0 | 0 |
| 2013 | 161 | **53** | 0 | 0 |
| 2014 | 242 | **61** | 0 | 0 |
| 2015 | 201 | **73** | 21 | 3 |
| 2016 | 272 | **62** | 55 | 28 |
| 2017 | 263 | **66** | 119 | 96 |
| 2018 | 223 | **65** | 154 | 116 |
| 2019 | 239 | **46** | 188 | 188 |

Surprisingly, a thorough comparison has yet to be made between TWAS and kernel methods. In their pioneering work on PrediXcan, Gamazon *et al.*[6] applied SKAT and PrediXcan to Wellcome Trust Case Control Consortium (WTCCC) data[21], reporting that PrediXcan produced an elevated proportion of significant genes across all P-values (Fig. 7 in [6]). However, the authors did not conduct simulations (under which the ground truth is known) to quantify the power of competing methods. Further developments to TWAS have incorporated multiple tissues[22], better models for predicting expression[9], methods to combating artifacts caused by co-expressed genes[23] and extensions to other middle-omes such as proteins[24, 25] and images[26]. These later developments have only been compared against the seminal TWAS tools[6, 8], and not directly with kernel methods such as the flagship tool, SKAT. Not only has this lack of comparisons unfairly discouraged the use of kernel methods, but there is also a missed opportunity for integrating the advantages of both approaches to better model the genetic basis of complex traits.

In this work, we propose a novel model called kTWAS (kernel-based transcriptome-wide association study), which integrates TWAS and kernel methods. We expect that kTWAS will take advantage of expression data via TWAS-based feature selection, and take advantage of the kernel-based test, which is robust to the unknown (possibly non-linear) underlying genetic architecture of the focal phenotype. As a result, the power of kTWAS should be equivalent to TWAS, due to its ability to select genetic variants regulating gene expressions; and also as robust as SKAT to noise and interactions between associated genetic variants.

Using simulated data, we have conducted thorough power comparisons between six protocols: PrediXcan, kTWAS, and four different protocols using SKAT under different assumptions regarding the distributions of genetic effects. (Detailed descriptions and justifications are presented in **Materials & Methods**). Although our main focus is on cases where subject-level genotypes are available, we have also tested six corresponding meta-analysis protocols where association mapping is conducted using summary statistics instead of subject-level genotypes. We simulate phenotypes based on four representative genetic architectures: an additive architecture, heterogeneous architecture, and two interaction architectures, with multiple effect sizes and heritability levels. While each protocol has unique strengths, our kTWAS method outperforms alternatives with significant margins in the majority of cases. As expected, the corresponding meta-analysis protocols had similar performance trends in our meta-analysis simulations. Moreover, we have conducted extensive real data analysis using kTWAS, which identified a larger number of significant genes with supporting literature than standard TWAS. Evidences from literature search support the real phenotypic validity of the novel genes discovered by kTWAS.

The following section presents the design of kTWAS, simulation details, and the power analysis procedure. The simulation outcomes and discoveries in real data are presented in Results. Finally, the conclusion discusses the potential impact of this work, additional literature, and future directions.

## Materials & Methods

### Mathematical details of SKAT, PrediXcan, and kTWAS

The sequence kernel association test (SKAT) tool was selected to represent kernel-based methods. SKAT utilizes a score test to aggregate the phenotypic contributions of multiple genetic variants using a kernel machine[20]. In particular, SKAT employs a score test:

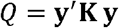

Where ***y*** is the vector of phenotype values, and ***K*** is a kernel calculated from the centralized genotype matrix ***G***, where *G_ij_* is the variant of the *j-*th genomic position in the GWAS focal region of the *i-*th individuals with. A simple example of a linear kernel is given by **K** = **G′(I)G**/n (where n is the total number of variants in the GWAS dataset).

While Wu *et al.*, originally subtracted cofactors such as sex and age from the phenotype vector **y** as **‘y-u’**[20], to simplify comparisons, this work will not include any cofactors when evaluating each model. Furthermore, while additional extensions to SKAT have been developed to handle rare variants[19, 27] and the combined effect of rare and common variants[28], this paper will only focus on common variants to be more comparable to TWAS.

As the earliest TWAS method, PrediXcan was selected to represent TWAS in this paper. PrediXcan is composed of two steps: First, a linear model is trained to predict genetically regulated gene expression (called GReX[6]) in the relevant tissue using a reference panel containing both genotype and expression data:

Here, are the regression parameters to be trained; and is the matrix of genotype in the focal region. Various methods can be used to train this predictive model[8, 29], and PrediXcan uses ElasticNet[30] to conduct this training. We use the pre-trained PrediXcan ElasticNet models which are available for download on the authors’ website[31].

Using the above model, GReX expressions are then estimated for the genotypes from GWAS dataset (which only provides genotype and phenotype information):

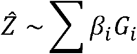

The estimated GReX values Ẑ, are then associated to the phenotype:

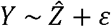

Various extensions to PrediXcan have since been developed for many cases in association mapping[8, 22, 23, 32, 33]. In particular, Gusev *et al.* pioneered the first tool utilizing summary statistics[8] to conduct TWAS. To ensure theoretical and technical consistency, in this paper we chose S-PrediXcan, the meta-analysis version of PrediXcan[12], to represent meta-analysis TWAS tools in our protocol comparisons.

Based on the hypothesis that SKAT and TWAS have different advantages which can be integrated, we developed the novel method kTWAS, or kernel-based transcriptome-wide association study. The protocol of kTWAS is illustrated in **Fig. 1**.

**Figure 1.**
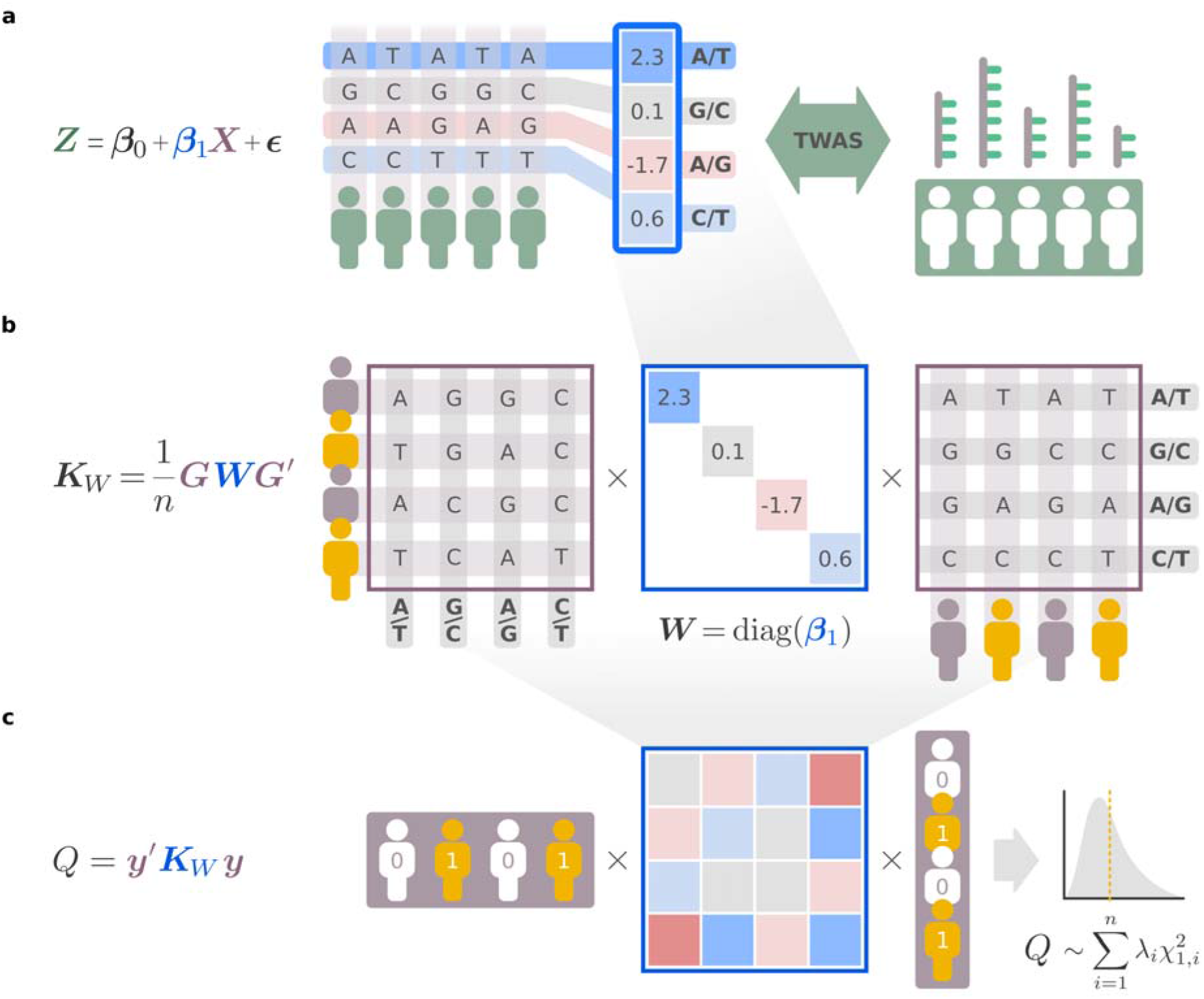
The kTWAS protocol. **(a)** A pretrained model (such as the ElasticNet model from PrediXcan) is used to linearly associate variants in a focal region to gene expressions, using genotype-expression data from a reference panel such as GTEx. **(b)** The regression parameters of the genotype-expression model from **a** are used to calculate the kernel for the sequence kernel association test (SKAT), where is based on the weight of each variant from the parameters of the pretrained linear model. **(c)** Using the TWAS-informed kernel from **b**, the Q score test from SKAT is conducted on the GWAS dataset to test the hypothesis that the variance components of a linear GWAS model are uniformly zero. Q follows a mixture of chi-squared distributions under the null hypothesis.

Mathematically, we first extract the regression coefficients *β*_*i*_ from the pretrained genotype-expression ElasticNet model provided by PrediXcan (**Fig. 1a**):

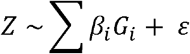

We then prepare the kernel ***K***_*W*_ for use in SKAT, where ***K***_*W*_ is weighted according to the contribution of each variant to the ElasticNet model above, ***W*** = *diag*(*β*_1_, … *β*_*m*_),(**Fig. 1b**):

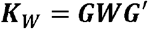

Finally, we conduct the Q score test from SKAT using the TWAS-informed kernel ***K***_*W*_. This tests the hypothesis that the variance components explained by the local genetic region are uniformly zero, where Q follows a mixture of chi-squared distributions under the null hypothesis (**Fig. 1c**):

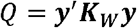

As outlined in the Introduction, kTWAS should enjoy the advantages of both kernel methods and TWAS, allowing it to incorporate both feature selection and feature modeling using a kernel.

### Protocols compared

We selected a total of six genotype-based protocols for power comparisons, including kTWAS.

1. SKAT-naive applies the default setting of SKAT, which does not select a subset of genetic variants in a region. In practice, researchers may use this “naive” version of SKAT given no prior knowledge of the relative importance of variants in the focal region.
2. SKAT-S-LM pre-selects genetic variants based on their marginal associations to phenotype, which is assessed by associating each individual variant in the region independently to the phenotype. A linear model, such as the model implemented in PLINK[34], is used to pre-select an arbitrary number of variants. As different genes have wildly varying numbers of causal genotypes contributing to the phenotype, we chose a pragmatic approach where the number of variants selected by SKAT-S-LM is matched to the number of variants selected by the ElasticNet model from PrediXcan.
3. SKAT-S-LMM is similar to SKAT-S-LM, but uses a linear mixed model (LMM) to perform variant selection, as implemented by EMMAX[35]. Since LM and LMM are both representative models for conducting single-variant GWAS, we chose to test both to cover a wider spectrum of variant selection methods.
4. SKAT-eQTL pre-selects genetic variants based on published eQTLs[36], instead of screening for marginal effects in the GWAS dataset under analysis. Since this protocol selects eQTLs independently of the GWAS data, it may allow SKAT-eQTL to avoid overfitting which may be caused by associating GWAS markers directly to phenotype, such as in the cases of SKAT-S-LM and SKAT-S-LMM. As such, we expect the performance of SKAT-eQTL to show different behavior from SKAT-S-LM and SKAT-S-LMM, depending on the marginal effects of individual variants. These eQTLs are downloaded from the GTEx publication[36], which are also selected by associating expressions to genotypes. The difference between this selection and the ElasticNet selection is that the variants selected are not jointly modeled linearly.
5. PrediXcan, as discussed previously, is the first and most representative TWAS tool.
6. kTWAS, which is our novel tool integrating TWAS and SKAT.

Additionally, we conducted an equivalent comparison of the above six protocols when applied to meta-analysis, based on the protocols MetaSKAT[27] and S-PrediXcan[12]. In all of the protocols that SKAT is relevant (including kTWAS), the default linear kernel is used in SKAT (https://cran.r-project.org/web/packages/SKAT/SKAT.pdf). Note that, as will be evidenced in the Results, a linear kernel is also more robust to non-linear effects than linear combinations adapted in TWAS. This is because the probability of two subjects carrying the same combinations of genetic variants is proportional to their genetic similarity captured by any (including linear) kernels.

### Data simulation procedure and power analysis

#### Genotype data and selected gene region

We used genotype data with a sample size of N = 2,548 from the 1000 Genomes Project[37] (available at http://hgdownload.cse.ucsc.edu/gbdb/hg38/1000Genomes/). We used the pretrained genotype-expression ElasticNet models used by PrediXcan[6, 12], available at http://predictdb.org/. The ElasticNet models are trained for all available tissue types in GTEx (v8). As the sample size and data quality varies between different GTEx tissues, the number of genes for which PrediXcan is applicable also varies. Whole-blood tissue is the largest and most frequently used gene set, with 7,252 available genes (together with 1Mb flanking genetic regions) on which PrediXcan can be applied. We therefore simulated 7,252 datasets based on the whole-blood tissue gene set.

#### Genetic architecture and parameterizations

For each gene, we simulated phenotypes based on four different genetic architectures. Their definitions and parameterizations are described below.

(1) “Additive” architecture. In this architecture, phenotype is associated with the sum of genetic effects. For each gene, we selected a genetic region that includes the gene body and 1Mb of flanking sequences. From this region, 4 single nucleotide polymorphisms (SNPs) with minor allele frequency (MAF) higher than 1% were randomly selected, with 2 SNPs chosen from variants preselected by the ElasticNet model used in PrediXcan, and the other 2 SNPs chosen from known eQTLs excluding those identified by the ElasticNet model. The first category of ElasticNet SNPs (selected by the genotype-expression ElasticNet model) favor the performance of PrediXcan, while the second category of eQTL SNPs (not selected by the ElasticNet model) favor SKAT-related models, as kernels better capture the effects of unsampled variants. To simplify simulations, we fix the number of SNPs from each category to 2, and we further rescale the phenotypic variance components contributed by ElasticNet SNPs versus other eQTL SNPs by a “scale” parameter which is set to one of six different factors: 0.0, 0.2, 0.4, 0.6, 0.8, and 1.0.
(2) “Heterogeneous” architecture. In this architecture, we randomly select two SNPs in the focal region. Subjects carrying an alternate allele at either or both SNPs will have an associated phenotypic change, where subjects carrying both alternate alleles have the same phenotypic change as subjects carrying one alternate allele. As in the additive model, the use of ElasticNet SNPs (favoring TWAS) versus eQTL SNPs (favoring kernel methods) is adjustable. We introduce a “proportion” parameter to set the number of ElasticNet SNPs as 0, 1, and 2, where the remaining number of eQTL SNPs is 2, 1, and 0 respectively. This parameter is analogous to the “scale” parameter applied to the variance components of ElasticNet SNPs in the additive architecture.
(3) & (4) “Recessive” and “Compensatory” interaction architectures. Similar to the heterogeneous architecture, we randomly select two SNPs in the focal region, and also include the “proportion” parameter which selects for 0, 1, and 2 ElasticNet SNPs. The effects of the SNPs are modeled differently, however. In the “Recessive” interaction architecture, a phenotypic change is made only when both alternate alleles are present. For the “Compensatory” interaction model, the phenotypic change is made only if there is exactly one alternate allele out of the two SNPs. Subjects carrying both alternate alleles will have the same phenotypic change as subjects carrying neither of the two alleles. This mirrors a situation where the effect of one mutation is compensated by the presence of another mutation, which is a phenomenon observed frequently in many organisms[38–41].

#### Heritability

The above genetic architectures define how genetic components of a phenotype could be specified. Using the genetic components, we generated phenotypes where the variance component of genetics, or heritability, equals a preselected value *h*^2^. That is, given the variance of the phenotype’s genetic component as 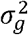, we calculate 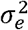 so that 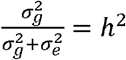. We then sample from the normal distribution 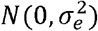 to determine the strength with which non-genetic components such as noise or environmental effects contribute to phenotype. Finally, the sum of the genetic and non-genetic components is stored as the simulated phenotype for use in association mapping and power calculations.

#### Power calculations & Adjustments to type-I errors

For each of the genetic architectures and their associated parameters, we simulated 7,252 datasets containing 2,548 subjects each, on different focal regions (containing a gene flanked by 1Mb sequences) in which causal variants are randomly selected. We then test each protocol’s ability to successfully identify the causal gene in each dataset, where success is defined as a Bonferroni-corrected[42] P-value that is lower than a predetermined critical value. We aimed to fix the type-I error across all protocols to α = 0.05. However, due to various reasons including the uneven distribution of genetic variants among the 7,252 genes and inherent biases between the protocols (e.g., overfitting caused by SKAT-S-LM and SKAT-S-LMM models), we discovered that the actual type-I errors of different protocols varies widely under a fixed critical value of 0.05. To equalize the type-I error across all protocols, we simulated random phenotypes with no genetic components whatsoever, to empirically determine the null distribution of each protocol. We then analyzed data from all 7,252 genetic regions using each protocol to empirically determine the critical value which separates out the smallest (most significant) 5% of all P-values. This ensures that all protocols are fairly compared with a type-I error of 0.05.

The statistical power of each protocol is given by the number of successes divided by the total number of datasets (7,252). For the six protocols utilizing genotype data, we conduct association mapping directly on the simulated genotypes. For the six protocols utilizing summary statistics, we conduct association mapping on summary statistics calculated from the simulated genotypes according to the instruction manuals of S-PrediXcan and MetaSKAT.

### Real data analysis

We compared the performance of kTWAS and PrediXcan on the WTCCC[43] and MSSNG[44] datasets. WTCCC contains 2,000 individual genotypes for each of the 7 complex diseases, of primarily European ancestry, along with 3,000 shared controls. The diseases surveyed by WTCCC are bipolar disease (BD), coronary artery disease (CAD), Crohn’s disease (CD), rheumatoid arthritis (RA), type 1 diabetes (T1D), type 2 diabetes (T2D), and hypertension (HT). Genotype data was collected from individuals using Affymetrix GeneChip 500K arrays. Following the PrediXcan paper, we used the whole-blood expressions to analyze all diseases. MSSNG is the largest available whole genome sequencing dataset for Autism Spectrum Disorder (ASD), containing 7065 sequences from ASD patients and controls[44]. As Cerebellum is reported as the most relevant tissue to ASD[45–47], we used its expression in GTEx data in the analysis.

## Results

### Simulations

#### Type-I errors & cutoffs

The 5% cutoff (determined by simulating the null distribution of each protocol) are generally close to the targeted type-I error of α = 0.05, except in the cases of SKAT-S-LM and SKAT-S-LMM (**Table 2**). This is consistent with the intuition that the pre-selection process in SKAT-S-LM and SKAT-S-LMM amplifies random false effects, which inflates type-I errors for these protocols. We apply a more stringent cutoff (determined from the simulations described above), to ensure fairness in the power comparisons below.

**Table 2.** Cutoffs that ensure Type-I error being 0.05 for compared protocols.

| Model | 5% cutoff | Model | 5% cutoff |
| --- | --- | --- | --- |
| SKAT-naive | 5.50E-02 | MetaSKAT-naive | 5.36E-02 |
| SKAT-eQTL | 5.22E-02 | MetaSKAT-eQTL | 5.42E-02 |
| SKAT-S-LM | 1.12E-14 | MetaSKAT-S-LM | 1.04E-14 |
| SKAT-S-LMM | 1.27E-14 | MetaSKAT-S-LMM | 1.22E-14 |
| PrediXcan | 5.04E-02 | S-PrediXcan | 5.79E-02 |
| kTWAS | 5.29E-02 | Meta-kTWAS | 4.97E-02 |

#### Additive architecture

**Fig. 2** and **Fig. 3** plot the power of genotype and summary statistic-based protocols under the additive model of genetic architecture. kTWAS clearly outperforms PrediXcan at all scale factors of the contribution from ElasticNet SNPs, showing that kernel methods using TWAS-based feature selection can always outperform the linear model utilized by TWAS, even when the underlying genetic architecture is also linear.

**Figure 2.**
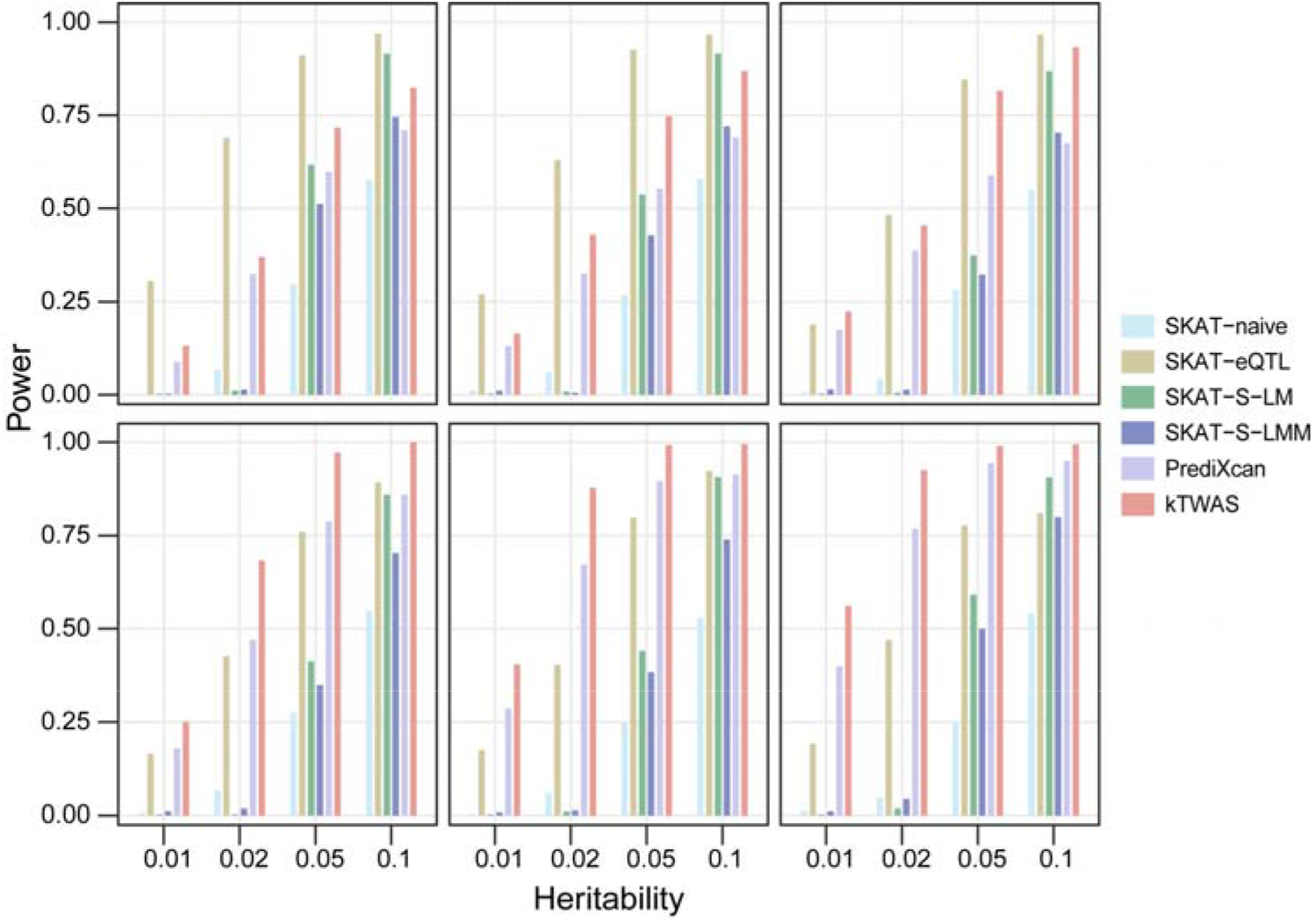
Statistical power (y-axis) of genotype-based protocols compared on the additive architecture at varying levels of trait heritability (x-axis) and contribution from ElasticNet SNPs. The compared protocols are SKAT-naive, SKAT-eQTL, SKAT-S-LM, SKAT-S-LMM, PrediXcan, and kTWAS. The scale factors applied to ElasticNet SNPs are 0.0, 0.2, 0.4 in the top row (left to right), and 0.6, 0.8, 1.0 in the bottom row (left to right).

**Figure 3.**
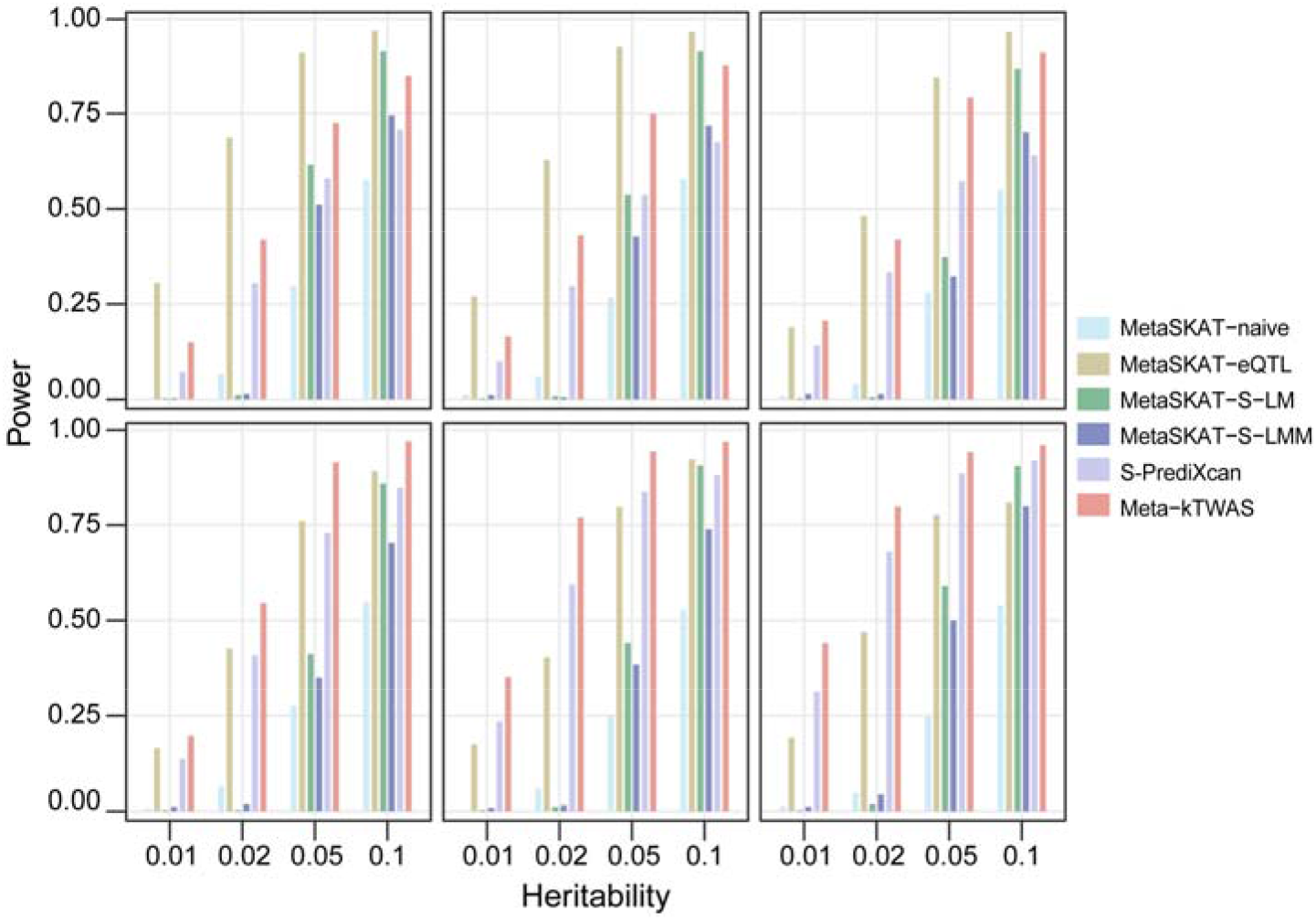
Statistical power (y-axis) of meta-analysis protocols compared on additive architecture at varying levels of trait heritability (x-axis) and contribution from ElasticNet SNPs. The compared protocols are MetaSKAT-naive, MetaSKAT-eQTL, MetaSKAT-S-LM, MetaSKAT-S-LMM, S-PrediXcan, and Meta-kTWAS. The scale factors applied to ElasticNet-selected SNPs are 0.0, 0.2, 0.4 in the top row (left to right), and 0.6, 0.8, 1.0 in the bottom row (left to right).

Comparisons between the four SKAT-based methods show that SKAT-eQTL performs best when the proportion of ElasticNet SNPs is low. This is expected since the eQTL SNPs (that are not in the ElasticNet selected list) favor kernel methods. When the proportion of ElasticNet SNPs is high, favoring the PrediXcan model, SKAT-eQTL has worse performance than both kTWAS and PrediXcan. SKAT-S-LM and SKAT-S-LMM, which select SNPs based on marginal effects, are generally less powerful than SKAT-eQTL and kTWAS, indicating that their pre-selection process may overfit the GWAS data and therefore reduce power (even after type-I errors are adjusted to be equivalent). When regional heritability is low, the power of SKAT-S-LM and SKAT-S-LMM are both extremely low, likely due to noise caused by random artifacts. Overall, SKAT-naive, which does not pre-select variants, has the lowest power when heritability is greater than 0.05, but outperforms SKAT-S-LM and SKAT-S-LMM when heritability is less than 0.05. In particular, at a very high heritability of *h*^2^ = 0.1, SKAT-S-LM and its meta-analysis equivalent have very high power approaching that of SKAT-eQTL and kTWAS. This may be because SNPs have strong marginal associations when overall genetic effects are high, which reduces noise in the linear pre-selection process employed by SKAT-S-LM and SKAT-S-LMM. We do not have a clear interpretation on why SKAT-S-LM consistently outperforms SKAT-S-LMM.

The meta-analysis protocols utilizing summary statistics exhibit similar performance compared to their genotype-based counterparts, although their overall power is slightly lower than that of genotype-based protocols.

#### Non-linear architectures

***Figs. 4***, ***5***, and ***6*** plot the power of genotype and summary statistic-based protocols under the Heterogeneous, Recessive interaction, and Compensatory interaction genetic architectures. Although these architectures fundamentally differ, several trends are consistent across all architectures: 1) kTWAS always outperforms PrediXcan; 2) SKAT-eQTL outperforms kTWAS when both causal SNPs are eQTL SNPs (that are not in the ElasticNet selected list); 3) SKAT-S-LM has high power only when heritability is high. Notably, under these non-linear architectures kTWAS and SKAT-eQTL outperform PrediXcan with larger margins than in the additive model. This is consistent with our expectation that kernel methods adapt better to the presence of genetic interactions, even when the kernel is linear.

**Figure 4.**
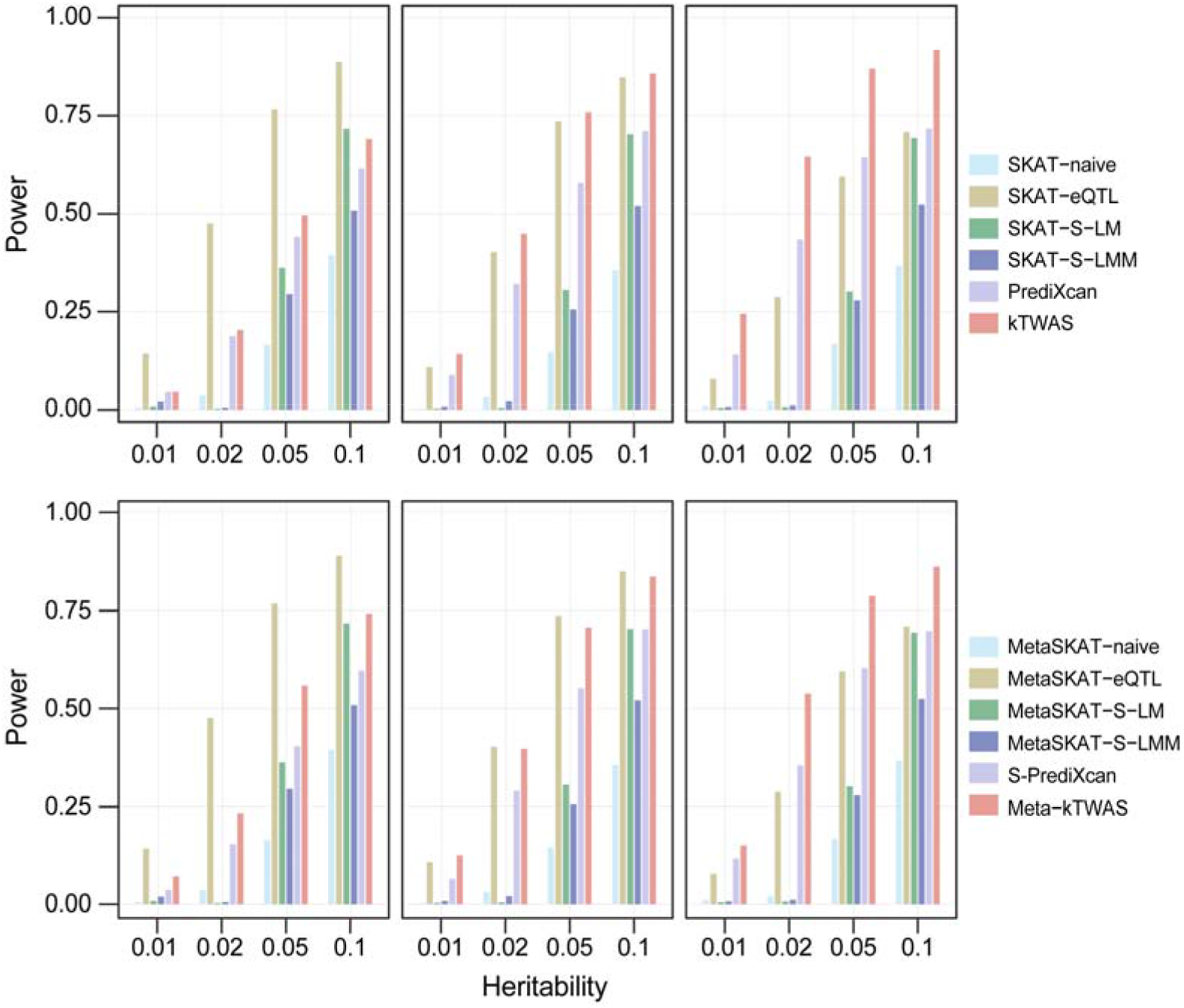
Statistical power (y-axis) of protocols of genotypes (top row) and summary statistics (bottom row), compared on Heterogeneous architecture at varying levels of trait heritability (x-axis) and different proportions of ElasticNet SNPs. The number of ElasticNet SNPs in both rows is 0, 1, 2 (left to right), and the corresponding number of eQTL-selected SNPs is 2, 1, 0.

**Figure 5.**
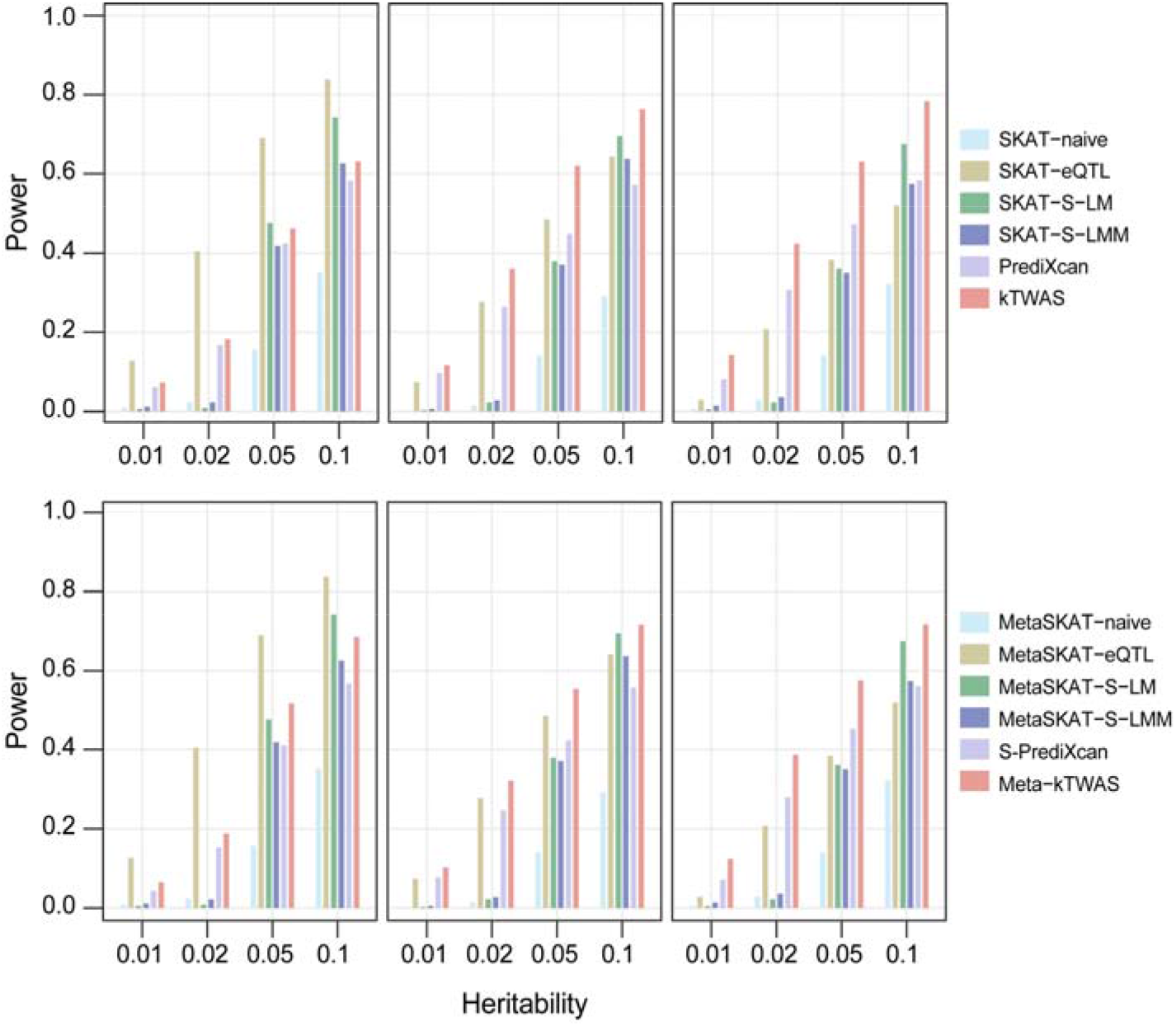
Statistical power (y-axis) of protocols of genotypes (top row) and summary statistics (bottom row) compared on Recessive architecture simulated at varying levels of trait heritability (x-axis) and different proportions of ElasticNet SNPs. The number of ElasticNet SNPs in both rows is 0, 1, 2 (left to right), and the corresponding number of eQTL SNPs is 2, 1, 0.

**Figure 6.**
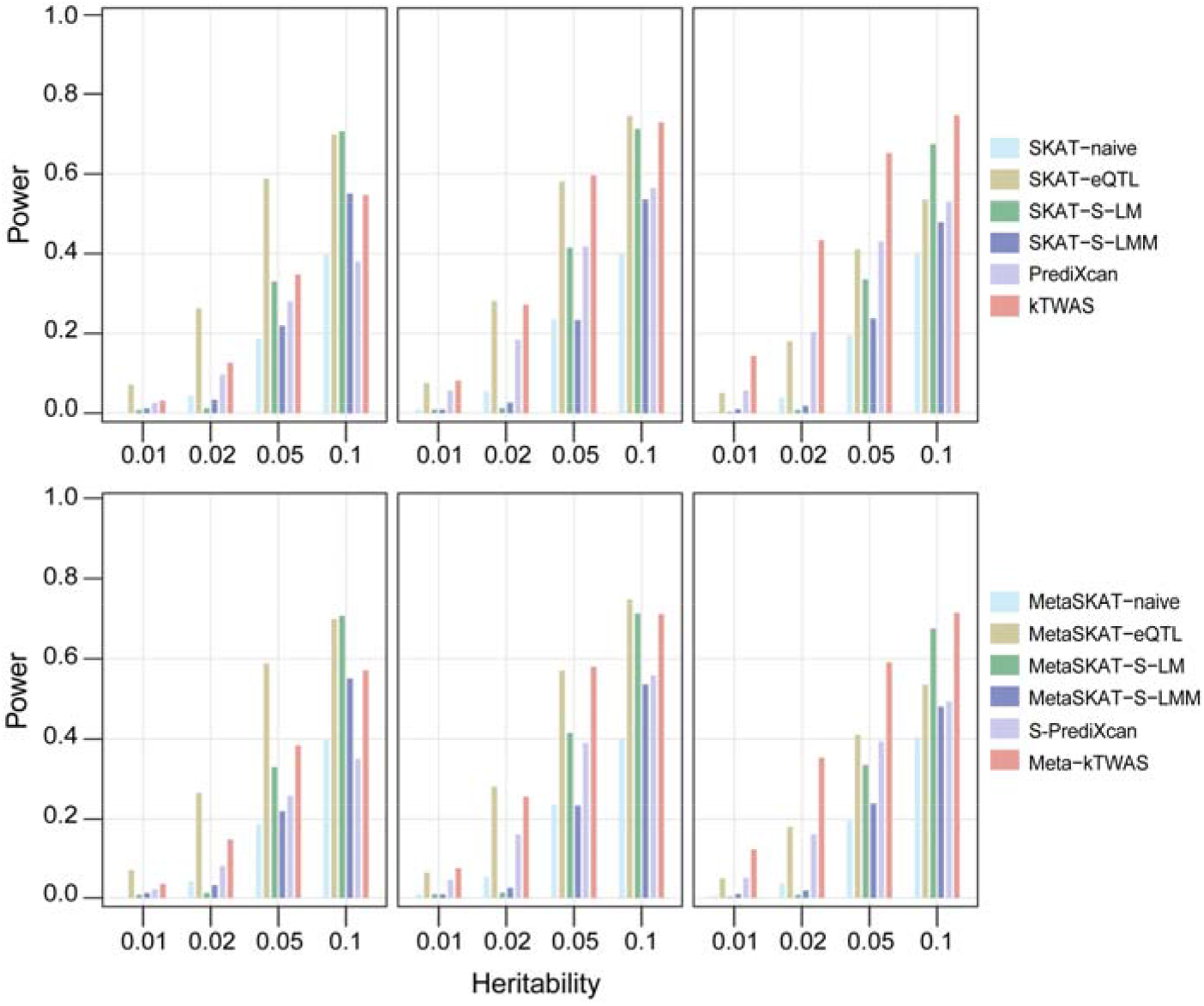
Statistical power (y-axis) of protocols of genotypes (top row) and summary statistics (bottom row), compared on Compensatory architecture simulated at varying levels of trait heritability (x-axis) and different proportions of ElasticNet SNPs. The number of ElasticNet SNPs in both rows is 0, 1, 2 (left to right), and the corresponding number of eQTL SNPs is 2, 1, 0.

### Applying kTWAS to real data

#### ASD whole genome data provided by MSSNG

**Fig. 7a** shows the Manhattan plot for the output of kTWAS. Based on a Bonferroni-corrected P-value < 0.05, we observed 6 peaks corresponding to RP11-575H3.1 (nominal P=1.73×10^−6^), NDUFV1 (P=2.06×10^−6^), PPP1R32 (P=2.59×10^−6^), NBPF15 (P=3.11×10^−6^), NBPF9 (P=5.82×10^−6^), and SRGAP2B (P=6.44×10^−6^). **Fig. 7b** shows the corresponding Manhattan plot for PrediXcan. Two genes (RP11-575H3.1 and NBPF15) identified by kTWAS were also discovered by PrediXcan, but at weaker significance levels (nominal P-values of 2.74×10^−6^ and 7.17×10^−6^, respectively). The remaining four genes are not identified as significant with PrediXcan (nominal P-values of 0.23 for SRGAP2B, 1.31×10-3 for NBPF9, 0.66 for NDUFV1, 4.67×10^−4^ for PPP1R32).

**Figure 7.**
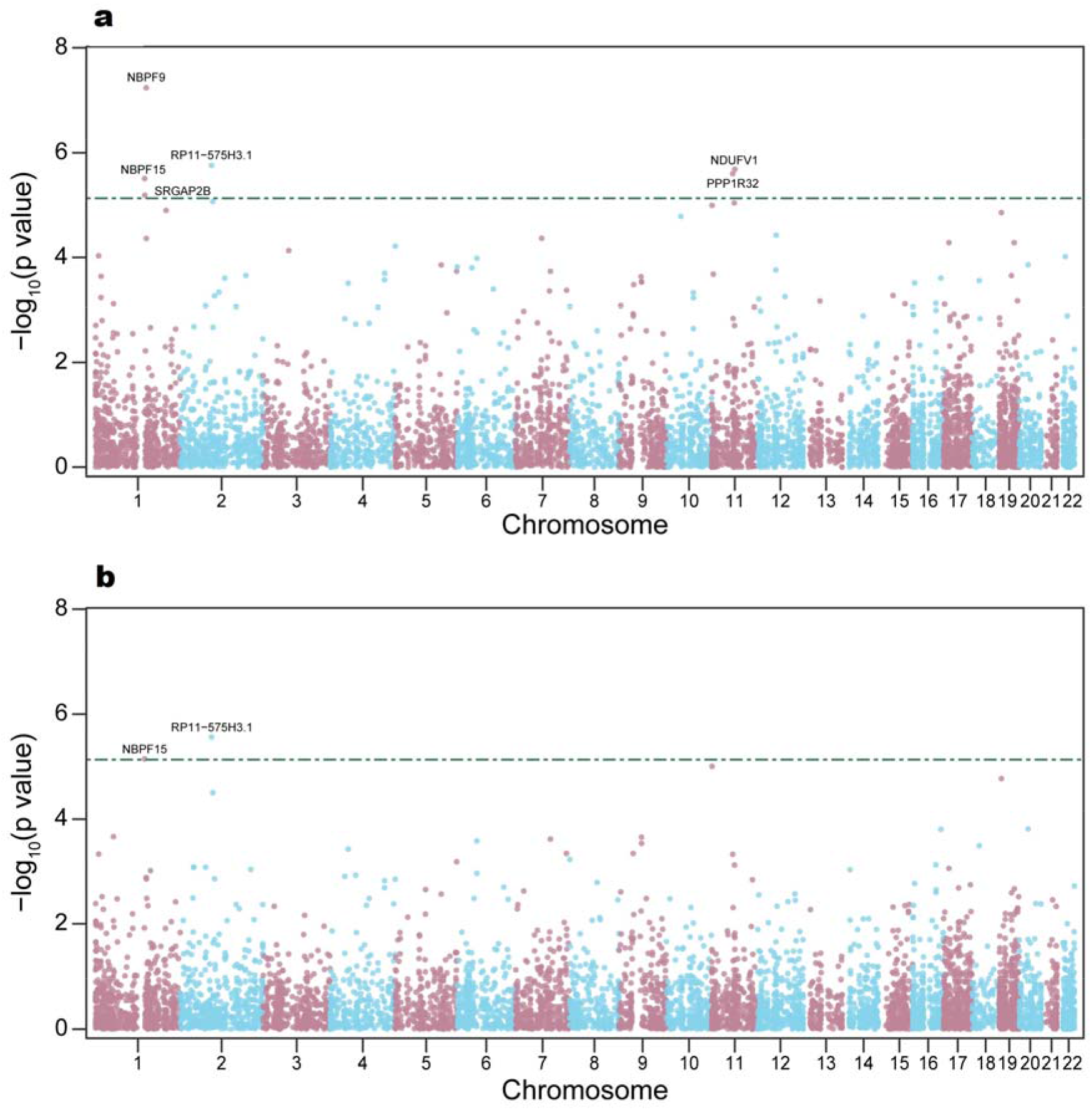
GWAS Manhattan plots of negative log P-values (y-axis) for Autism-associated SNPs at genome coordinates (x-axis) of MSSNG consortium whole genome data. Plots show associations generated by kTWAS (**a**) and PrediXcan (**b**), with gene expressions predicted using the GTEx cerebellum ElasticNet model. The Bonferroni-corrected significance threshold (green dashed line) is 7.37×10^−6^ (= 0.05/6794).

Out of the four genes identified only by kTWAS, three have literature supporting their association with ASD. The inhibition of SRGAP2 function by its human-specific paralogs has contributed to the evolution of the human neocortex and plays an important role during human brain development[48, 49]. NBPF9 is a member of the neuroblastoma breakpoint family (NBPF) which consists of dozens of recently duplicated genes primarily located in segmental duplications on human chromosome 1. Members of this gene family are characterized by tandemly repeated copies of DUF1220 protein domains. Gene copy number variations in the human chromosomal region 1q21.1, where most DUF1220 domains are located, have been implicated in a number of developmental and neurogenetic diseases such as autism[50]. In particular, rare variants located in NBPF9 are reported to be associated with ASD[51]. Additionally, evidence shows that NDUFV1 is a ‘developmental/neuropsychiatric’ susceptibility gene when a rare duplication CNV occurs at 11p13.3[51]. The only gene not supported by literature is PPP1R32, which may be a novel gene for ASD research. Both genes identified jointly by kTWAS and PrediXcan are supported by literature[52, 53].

#### WTCCC genotyping data

We applied kTWAS to type 1 diabetes (T1D) data, identifying 52 genes significantly associated with risk of T1D (Bonferroni-corrected p-value < 0.05). In contrast, PrediXcan identified 32 genes, of which 31 were also detected by kTWAS (**Table 3**). Among the 21 genes identified only by kTWAS, 19 are within the MHC region which has been shown to influence susceptibility to complex, autoimmune, and infectious diseases including T1D in particular[54]. Most (except for four) of these genes have been reported as having associations with T1D[43, 55–68].

**Table 3.**
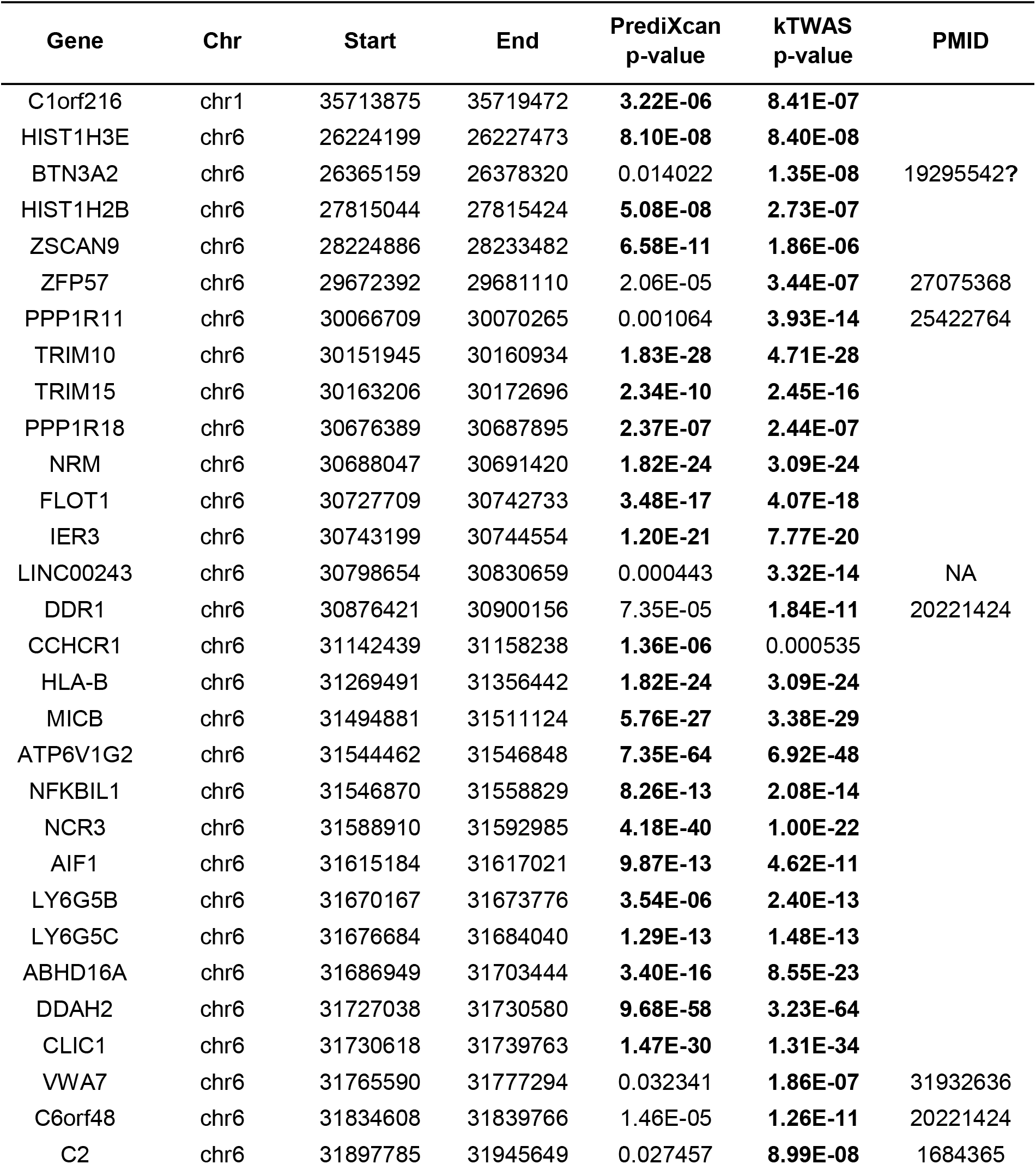
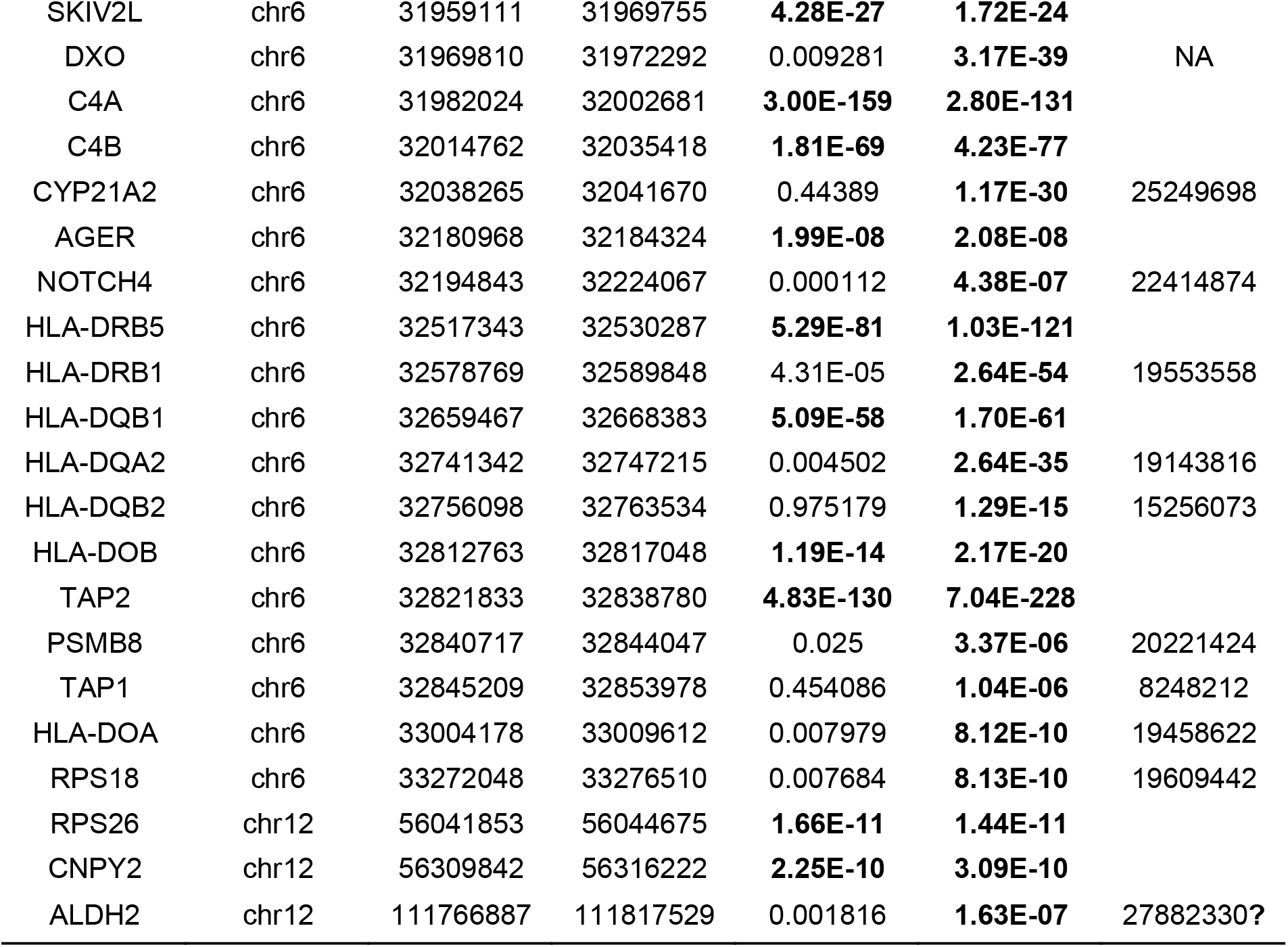
PrediXcan and kTWAS results for Bonferroni-corrected significant gene associations with type 1 diabetes in WTCCC data. To account for multiple testing, we used a significance threshold of 6.89×10^−6^ (0.05/7252) for all diseases. Significant genes are in bold. Chromosome and gene start positions are based on GENCODE version 26. The question marked PMIDs indicate relevant, however not supportive, literature.

| Gene | Chr | Start | End | PrediXcan<br>p-value | kTWAS<br>p-value | PMID |
| --- | --- | --- | --- | --- | --- | --- |
| C1orf216 | chr1 | 35713875 | 35719472 | **3.22E-06** | **8.41E-07** |  |
| HIST1H3E | chr6 | 26224199 | 26227473 | **8.10E-08** | **8.40E-08** |  |
| BTN3A2 | chr6 | 26365159 | 26378320 | 0.014022 | **1.35E-08** | 19295542? |
| HIST1H2B | chr6 | 27815044 | 27815424 | **5.08E-08** | **2.73E-07** |  |
| ZSCAN9 | chr6 | 28224886 | 28233482 | **6.58E-11** | **1.86E-06** |  |
| ZFP57 | chr6 | 29672392 | 29681110 | 2.06E-05 | **3.44E-07** | 27075368 |
| PPP1R11 | chr6 | 30066709 | 30070265 | 0.001064 | **3.93E-14** | 25422764 |
| TRIM10 | chr6 | 30151945 | 30160934 | **1.83E-28** | **4.71E-28** |  |
| TRIM15 | chr6 | 30163206 | 30172696 | **2.34E-10** | **2.45E-16** |  |
| PPP1R18 | chr6 | 30676389 | 30687895 | **2.37E-07** | **2.44E-07** |  |
| NRM | chr6 | 30688047 | 30691420 | **1.82E-24** | **3.09E-24** |  |
| FLOT1 | chr6 | 30727709 | 30742733 | **3.48E-17** | **4.07E-18** |  |
| IER3 | chr6 | 30743199 | 30744554 | **1.20E-21** | **7.77E-20** |  |
| LINC00243 | chr6 | 30798654 | 30830659 | 0.000443 | **3.32E-14** | NA |
| DDR1 | chr6 | 30876421 | 30900156 | 7.35E-05 | **1.84E-11** | 20221424 |
| CCHCR1 | chr6 | 31142439 | 31158238 | **1.36E-06** | 0.000535 |  |
| HLA-B | chr6 | 31269491 | 31356442 | **1.82E-24** | **3.09E-24** |  |
| MICB | chr6 | 31494881 | 31511124 | **5.76E-27** | **3.38E-29** |  |
| ATP6V1G2 | chr6 | 31544462 | 31546848 | **7.35E-64** | **6.92E-48** |  |
| NFKBIL1 | chr6 | 31546870 | 31558829 | **8.26E-13** | **2.08E-14** |  |
| NCR3 | chr6 | 31588910 | 31592985 | **4.18E-40** | **1.00E-22** |  |
| AIF1 | chr6 | 31615184 | 31617021 | **9.87E-13** | **4.62E-11** |  |
| LY6G5B | chr6 | 31670167 | 31673776 | **3.54E-06** | **2.40E-13** |  |
| LY6G5C | chr6 | 31676684 | 31684040 | **1.29E-13** | **1.48E-13** |  |
| ABHD16A | chr6 | 31686949 | 31703444 | **3.40E-16** | **8.55E-23** |  |
| DDAH2 | chr6 | 31727038 | 31730580 | **9.68E-58** | **3.23E-64** |  |
| CLIC1 | chr6 | 31730618 | 31739763 | **1.47E-30** | **1.31E-34** |  |
| VWA7 | chr6 | 31765590 | 31777294 | 0.032341 | **1.86E-07** | 31932636 |
| C6orf48 | chr6 | 31834608 | 31839766 | 1.46E-05 | **1.26E-11** | 20221424 |
| C2 | chr6 | 31897785 | 31945649 | 0.027457 | **8.99E-08** | 1684365 |
| SKIV2L | chr6 | 31959111 | 31969755 | **4.28E-27** | **1.72E-24** |  |
| DXO | chr6 | 31969810 | 31972292 | 0.009281 | **3.17E-39** | NA |
| C4A | chr6 | 31982024 | 32002681 | **3.00E-159** | **2.80E-131** |  |
| C4B | chr6 | 32014762 | 32035418 | **1.81E-69** | **4.23E-77** |  |
| CYP21A2 | chr6 | 32038265 | 32041670 | 0.44389 | **1.17E-30** | 25249698 |
| AGER | chr6 | 32180968 | 32184324 | **1.99E-08** | **2.08E-08** |  |
| NOTCH4 | chr6 | 32194843 | 32224067 | 0.000112 | **4.38E-07** | 22414874 |
| HLA-DRB5 | chr6 | 32517343 | 32530287 | **5.29E-81** | **1.03E-121** |  |
| HLA-DRB1 | chr6 | 32578769 | 32589848 | 4.31E-05 | **2.64E-54** | 19553558 |
| HLA-DQB1 | chr6 | 32659467 | 32668383 | **5.09E-58** | **1.70E-61** |  |
| HLA-DQA2 | chr6 | 32741342 | 32747215 | 0.004502 | **2.64E-35** | 19143816 |
| HLA-DQB2 | chr6 | 32756098 | 32763534 | 0.975179 | **1.29E-15** | 15256073 |
| HLA-DOB | chr6 | 32812763 | 32817048 | **1.19E-14** | **2.17E-20** |  |
| TAP2 | chr6 | 32821833 | 32838780 | **4.83E-130** | **7.04E-228** |  |
| PSMB8 | chr6 | 32840717 | 32844047 | 0.025 | **3.37E-06** | 20221424 |
| TAP1 | chr6 | 32845209 | 32853978 | 0.454086 | **1.04E-06** | 8248212 |
| HLA-DOA | chr6 | 33004178 | 33009612 | 0.007979 | **8.12E-10** | 19458622 |
| RPS18 | chr6 | 33272048 | 33276510 | 0.007684 | **8.13E-10** | 19609442 |
| RPS26 | chr12 | 56041853 | 56044675 | **1.66E-11** | **1.44E-11** |  |
| CNPY2 | chr12 | 56309842 | 56316222 | **2.25E-10** | **3.09E-10** |  |
| ALDH2 | chr12 | 111766887 | 111817529 | 0.001816 | **1.63E-07** | 27882330? |

The PMIDs of supporting literatures are listed in **Table 3**. The remaining four genes lacking literature support are BTN3A2, ALDH2, LINC00243 and DXO. Of these four novel genes, BTN3A2 is reported to play important roles in regulating the immune response, and is a potential novel susceptibility gene for T1D[55]. ALDH2 is known to offer myocardial protection against stress conditions such as diabetes mellitus[69], although the underlying mechanism is unclear.

The other diseases in WTCCC have limited numbers of significant genes, except in the case of rheumatoid arthritis (RA). kTWAS identified 24 genes associated with RA, while PrediXcan identified 19 significant genes, of which 18 are also detected by kTWAS (**Table 4**). All six genes identified only by kTWAS (VARS2, NCR3, NOTCH4, TAP2, HLA-DQB2, LY6G5B) are in the MHC region and have substantial literature support. In particular, a nonsynonymous change in the VARS2 locus (rs4678) is strongly associated with RA[70]. One SNP in NCR3 can regulate the expression of two genes in RA cases, and increased NCR3 expression is significantly associated with reduced RA susceptibility[71]. NOTCH4 is also reported to be RA-susceptible by multiple researchers[72, 73]. Yu *et al.* provided genetic evidence that TAP2 gene codon 565 polymorphism could play a role in RA[74]. A study on Italian patients found a mutation in HLA-DQA2 (rs9275595) could contribute to RA pathogenesis. Although there is no direct evidence to show LY6G5B is associated with RA, strong associations have been found between RA and a 126-kb region in the MHC class III region between BAT2 and CLIC1, which contains the five Ly-6 members including LY6G5B[75], indicating that LY6G5B might be a novel RA risk gene.

**Table 4.**
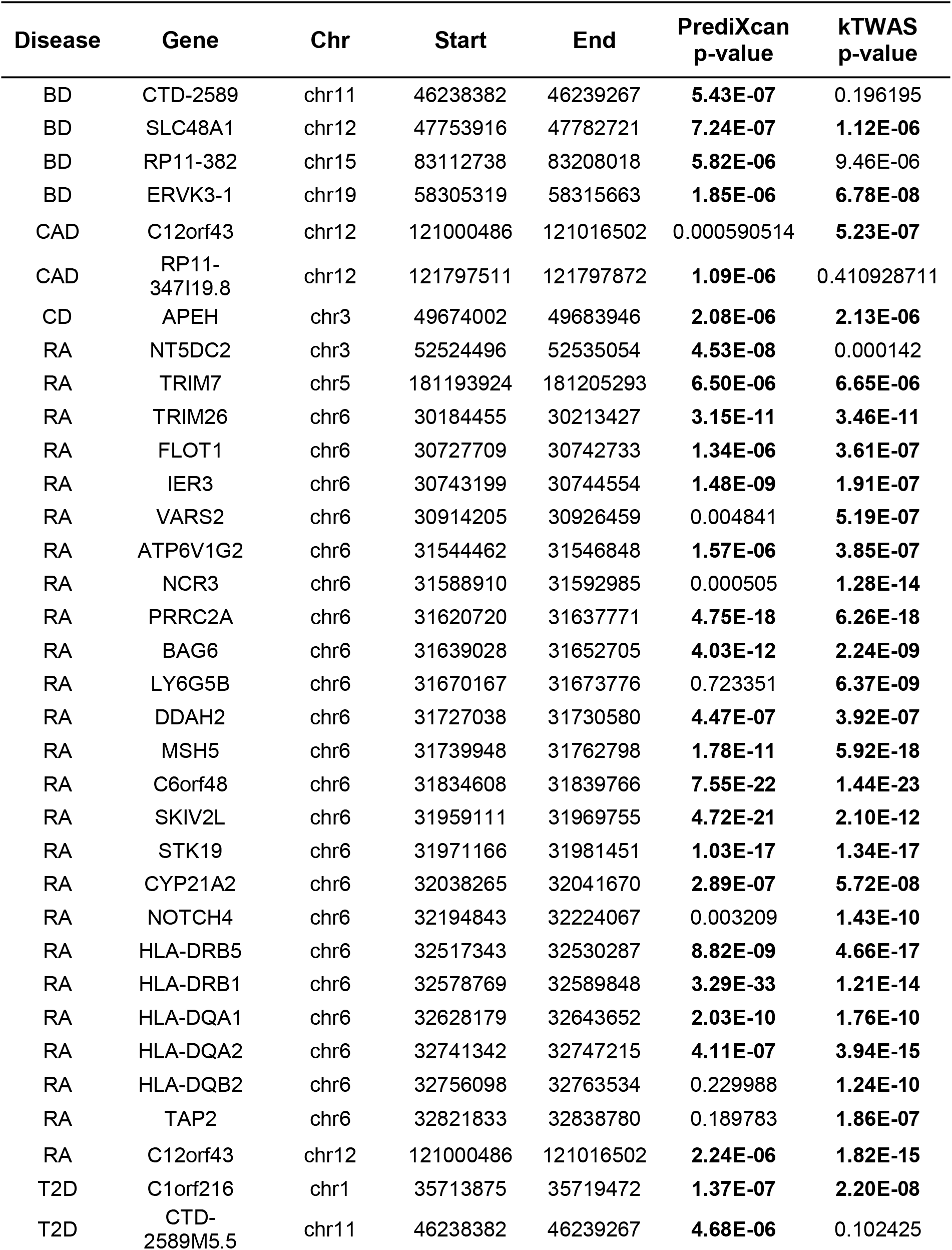
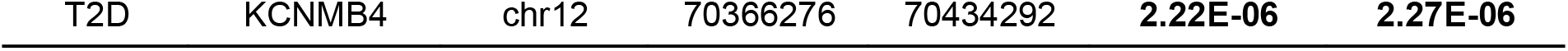
PrediXcan and kTWAS results for Bonferroni-corrected significant gene associations with five diseases in WTCCC consortium. To account for multiple testing, we used a significance threshold of 6.89×10^−6^ (0.05/7252) for all diseases. bipolar disease (BD), coronary artery disease (CAD), Crohn’s disease (CD), rheumatoid arthritis (RA), type 2 diabetes (T2D). The significant genes are in bold. Chromosome and gene start positions are based on GENCODE version 26.

Taken together, the above analyses of MSSNG sequence data and WTCCC genotype array data illustrate that kTWAS is able to identify a larger number of significant and meaningful genes in comparison to PrediXcan. These results confirm that the inclusion of kernel methods in TWAS increases statistical power in real and simulated data, whereas the use of linear combinations of selected SNPs in standard TWAS is unable to robustly model non-linear effects.

## Conclusion

In this work, we have thoroughly highlighted the essential advantages and differences between TWAS and kernel methods in terms of their ability to select and model genetic features. From this perspective, we have designed kTWAS, a novel protocol integrating the advantages of both methods in order to utilize expression data while being robust to non-linear effects. We demonstrate that kTWAS improves the power of TWAS, by conducting extensive simulations and real data analyses. This work will help researchers understand the ideal conditions for applying TWAS versus kernel methods, and provide a method which integrates them to capture non-linear effects. This work also reveals that linear kernels are more effective than simple linear regression for detecting non-linear genetic effects.

Other researchers have also investigated the link between SKAT and TWAS. Xu *et al.* have designed a power testing framework, where TWAS and SKAT are special cases of their test[29]. However, their framework does not directly compare the power of the two protocols, and they do not suggest a method for integrating the protocols.

As shown in **Figs. 2–6**, it is evident that SKAT-eQTL also has high power, despite not taking advantage of the TWAS-like feature pre-selection employed by ElasticNet or the multiple-regression based methods found in SKAT-S-LM and SKAT-S-LMM. In essence, SKAT-eQTL only selects for genetic variants with good marginal effects, and does not consider linear combinations of variants during feature selection. Our future work will thoroughly investigate the theoretical and experimental effectiveness of SKAT-eQTL via simulations and real data analyses.

### Key points

- New insights into TWAS and kernel methods are revealed. TWAS pre-selects and weights features in a linear model via expressions, whereas kernel methods conduct association analyses by modeling genetic similarity via various kernels. From the perspective of machine learning, these two methods cover two complementary aspects of feature engineering: feature selection/pruning, and feature modeling.
- A novel protocol called kTWAS is proposed, integrating transcriptome-wide association studies (TWAS) and sequence kernel association test (SKAT). Thorough testing shows this novel protocol enjoys the advantages of both TWAS and kernel-based models, resulting in increased power while being robust to non-linear effects.
- Twelve protocols based on TWAS and SKAT are thoroughly tested with four genetic architectures, under different heritability levels and other parameterizations.
- Novel genes are disclosed by applying kTWAS to WTCCC genotyping array data (seven diseases) and MSSNG sequence data (Autism Spectrum Disorder). kTWAS identified more significant genes with literature support than the competing TWAS protocol PrediXcan.

## Acknowledgement

Q.L. is supported by an NSERC Discovery Grant (RGPIN-2017-04860), a Canada Foundation for Innovation JELF grant (36605), a New Frontiers in Research Fund (NFRFE-2018-00748) and an ACHRI Startup grant. C.C. is supported by an ACHRI scholarship. D.K. is supported by an NSERC USRA award. S.E. is supported by an AIHS award.

